# Chronic alcohol intake regulates expression of SARS-CoV2 infection-relevant genes in an organ-specific manner

**DOI:** 10.1101/2022.02.01.478685

**Authors:** Marion M. Friske, Francesco Giannone, Mona Senger, Robin Seitz, Anita C. Hansson, Rainer Spanagel

**Affiliations:** Institute of Psychopharmacology, Central Institute of Mental Health, Heidelberg University, Germany

**Keywords:** COVID19, Ace2, Tmprss2, Mas, Alcohol Use Disorder

## Abstract

Chronic alcohol consumption and alcohol use disorder (AUD) have a tremendous impact on the patient’s psychological and physiological health. There is some evidence that chronic alcohol consumption influences SARS-CoV2 infection risk, but the molecular mechanism is unknown. Here, we generated expression data of SARS-CoV2 infection relevant genes (Ace2, Tmprss2 and Mas) in different organs in rat models of chronic alcohol exposure and alcohol dependence. ACE2 and TMPRSS2 represent the virus entry point whereas Mas is activating the anti-inflammatory response once the cells are infected. Across three different chronic alcohol test conditions, we found a consistent upregulation of Ace2 in the lung, which is the most affected organ in Covid-19 patients. Other organs such as liver, ileum, kidney, heart, and the brain showed also up-regulation of Ace2 and Mas but in a less consistent manner across the different animal models, while Tmprss2 was unaffected in all conditions. We suggest that alcohol-induced up-regulation of Ace2 can lead to an elevated stochastic probability of cellular virus entry and may thus confer a molecular risk factor for a SARS-CoV2 infection.

## Introduction

Since the end of 2019, the Coronavirus disease (COVID-19) has become a central topic of everyday life. The most relevant clinical syndrome of COVID-19 patients is impairment of the respiratory tract. Other organs, such as heart, kidney, liver, and the gastrointestinal tract are also affected by the virus (Li et al., 2020; Puelles et al., 2020; Xiao et al., 2020; Zou et al., 2020). In severely affected patients, neurological changes can be observed as well (Ellul et al., 2020; Liu et al., 2021a).

The mechanism of cell invasion by SARS-CoV2 has been studied extensively since the beginning of the pandemic and is now well understood. Once the virus enters its host via the mucous membranes of the nasal cave, it binds to the angiotensin converting enzyme 2 (ACE2) at the cell surface. With support of transmembrane protease serine subtype 2 (TMPRSS2), that cleaves the S1 unit of the viral spike protein, SARS-CoV2 is able to enter via cellular internalization and infect the cell (Hoffmann et al., 2020; Walls et al., 2020; Wrapp et al., 2020). Because of the internalization process, ACE2 concentration decreases on the outer cell membrane. In response to viral cell entry, ACE2 cleaves Angiotensin I to Angiotensin (1-9) and Angiotensin II to Angiotensin (1-7). As part of a protective mechanism, Angiotensin (1-7) binds primarily to the G-protein coupled receptor Mas on the cell membrane, which exerts anti-proliferative and anti-inflammatory effects, but also causes vasodilation, natriuresis, and diuresis (Kuriakose et al., 2021; Wiese et al., 2020).

More than 2 billion people drink alcohol regularly and alcohol has been classified as one of the most harmful drugs (Nutt et al., 2010). Chronic alcohol consumption can lead to alcohol dependence and caused 2.07 million deaths in males and 0.374 million deaths in females in 2019 (Murray et al., 2020). Chronic alcohol consumption and dependence can cause an impaired immune system, liver cirrhosis, cancer and complications associated with the respiratory system such as acute respiratory distress syndrome (ARDS) (Bailey et al., 2021; Corrao et al., 2004; Seitz et al., 2018; Simou et al., 2018; Zareini et al., 2021). Based on these harmful effects to the organism caused by chronic alcohol consumption, an elevated risk for severe COVID-19 infection and a worsened disease progression was hypothesized (Dubey et al., 2020; Mallet et al., 2020; Muhammad et al., 2021; Testino et al., 2022; Vasudeva & Patel, 2020). However, two studies reported that chronic alcohol consumption was not associated with an increased risk of COVID-19 infection or severe course of illness, respectively (Dai et al., 2020; Zhong et al., 2021). Interestingly, one group reported a positive association between alcohol consumption and COVID-19 infection until they corrected for closely related comorbid behaviors such as smoking, after which the association failed to stay significant (Hamer et al., 2020). This might hint to a secondary impact of alcohol when present in combination with smoking, as smoking has so far been shown by several studies to be a risk factor for COVID-19 infection and severe illness progression (Dorjee et al., 2020; Patanavanich & Glantz, 2021; Vardavas & Nikitara, 2020; Zheng et al., 2020). Although the epidemiological data do not give a definite answer of whether chronic alcohol consumption and dependence are risk factors for COVID-19 infection and disease severity, there might be a molecular link between alcohol and COVID-19 infection (Liu et al., 2021b).

To provide molecular insights into the interaction of chronic alcohol consumption and dependence on COVID-19 infection risk, we designed a study comparing RNA expression levels of some of the SARS-CoV2 infection-relevant genes including Ace2, Tmprss2 and Mas. We examined different organs that are usually affected by the virus, namely the lung, liver, small intestine (Ileum), kidney, heart, and the brain by using a standard protocol of subchronic repeated intermittent ethanol injections in rats over two weeks (Hundt et al., 1998).

We also used a well-established model of alcohol dependence, in which rats receive chronic intermittent exposure to alcohol vapor over 7 weeks (Meinhardt & Sommer, 2014). This leads to intoxication levels similar to those seen in clinical cases of Alcohol Use Disorder (AUD), and induces long-lasting behavioral as well as pronounced molecular changes in the brain. Most importantly, alcohol-dependent rats in this model show tolerance, persistent escalation of alcohol drinking, increased motivation to obtain alcohol, reduced cognitive flexibility, and increased relapse behavior (Meinhardt et al., 2021a; Meinhardt & Sommer, 2014; Rimondini et al., 2002; Sommer et al., 2008).

In summary, we hypothesize that chronic alcohol consumption and alcohol dependence lead to a change in expression of Ace2 and additional genes of the renin-angiotensinogen pathway involved in the SARS-CoV2 infection cascade. This effect might be organ specific and may constitute a molecular risk factor for COVID-19 infection and severe disease progression.

## Results

### Chronic alcohol exposure leads to an up-regulation of Ace2 in the lung

Since SARS-CoV2 binds to ACE2 to facilitate cell entry (Cuervo & Grandvaux, 2020), we first were interested in potential gene-expression changes of Ace2 caused by chronic alcohol consumption. Therefore, we used three different chronic ethanol treatment procedures: repeated intermittent ethanol intraperitoneal (IP) injections (sub-chronic treatment), vapor exposure for seven weeks (non-abstinence), and the post-dependent model (abstinence) (Meinhardt & Sommer, 2014). In the sub-chronic treatment procedure, animals received for two weeks a moderate dose of 1.5 g/kg IP twice per day with 6 h between the injections, with the first injection performed 2 hours after the beginning of the active cycle (Fig. 4). At the end of the experiment, we collected different organs that have been found to be potentially infected by SARS-CoV2 and that are known to express Ace2 at detectable levels. A factorial ANOVA, that was conducted to analyze organ-specific changes in Ace2 expression levels showed a significant main effect of Group (F[1, 48]=5.157, p=.028) indicating a higher Ace2 gene expression in the sub-chronic group compared to the control condition, and a significant interaction (F[4, 48]=3.065, p=.025). A post-hoc analysis revealed a significant effect in liver (p=.004, d=1.57) and lung (p=.006, d=2.32) with a more than two-fold up-regulation in the treatment group (Fig. 1A).

**Fig. 1:**
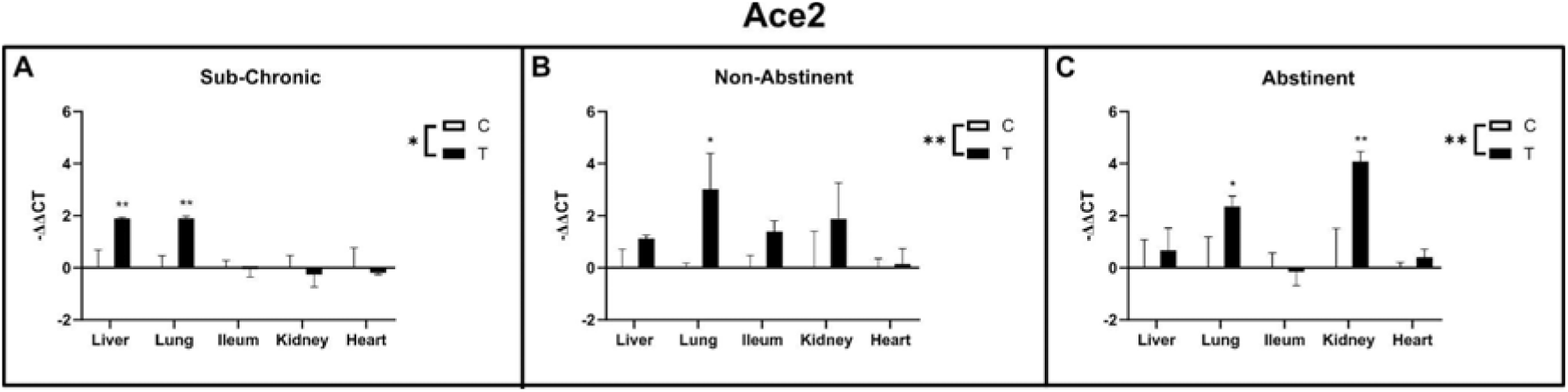
Ace2 gene expression in liver, lung, ileum, kidney and heart of sub-chronic ethanol IP injected (**A**) and ethanol vapor exposed animals (**B** non-abstinent, **C** abstinent) with respective control groups. Data are shown as mean ± SE. Factorial ANOVA shows significant differences between treatments (indicated as “T”) and control groups (indicated as “C”) for all three experiments. Subsequent post-hoc tests identified lung as significantly up-regulated in the treatment groups of all three experiments, whereas in the sub-chronic group liver and in the vapor abstinence group kidney was additionally up-regulated.

Next, a different batch of animals was exposed to ethanol vapor in a chronic intermittent ethanol treatment for seven weeks, 16 hours per day. Half of the batch was sacrificed immediately at the end of the last exposure cycle (non-abstinence group), the remaining half after three weeks of abstinence (abstinence/post-dependent group) (Fig. 5). For gene expression analysis, the same organs as for the sub-chronic treatment were collected. For both the non-abstinence and abstinence conditions we found a main effect of Group (non-abstinence: F[1, 43]=7.440, p=.009; abstinence: F[1, 49]=7.926, p=.007) with a significantly higher expression in the treated groups compared to their respective controls (Fig. 1B-C) and no interaction (non-abstinence: F[4, 43]=0.718, p=.585; abstinence: F[4, 49]=2.120, p=.092). Interestingly, in both of these treatments the post-hoc test revealed a significant upregulation of Ace2 in the lung (non-abstinence: p=.02, d=1.36; abstinence: p=.046, d=1.10), with the kidney also showing an upregulation in the abstinence condition (p=.002, d=1.52; Fig. 1B-C). Such consistent findings suggest that the lung might be more affected by chronic alcohol compared to the other organs. The post-hoc test resulted in a significant effect in lung (p=.046) and kidney (p=.002). Ileum and heart were unaffected regardless of treatment. (Fig. 1C).

In summary, Ace2 gene-expression was significantly up-regulated in all three procedures of chronic alcohol exposure. In the sub-chronic condition, the up-regulation was organ specific, with only lung and liver being significantly affected. Interestingly, the lung appeared to consistently show Ace2 upregulation in all conditions, as Ace2 mRNA levels were found to be significantly higher also in the alcohol vapor-exposed animals with and without abstinence compared to their respective controls (in the absence of a significant interaction) based on the post-hoc analysis. These results suggest that the lung might be more affected by chronic alcohol intake compared to the remaining organs observed. Particularly, Ace2 expression was found to be consistently increased in the lung, regardless of ethanol treatment regimen. Additionally, for the intermittent sub-chronic treatment procedure, the liver showed Ace2 upregulation which was also the case for the kidney following vapor exposure and abstinence. These experiments show an organ-specific up-regulation of Ace2, which might lead to a higher abundance of ACE2 protein and therefore an increased stochastic probability of the SARS-CoV2 to enter and infect the cells.

### Augmented anti-inflammatory response via the ACE2/Ang(1-7)/Mas cascade following alcohol abstinence

To enlarge the scope of our study, we additionally observed gene expression levels of Tmprss2 and Mas. TMPRSS2 functions as a co-player of ACE2 to enable SARS-CoV2 cell entry. Once the virus affects the cell, Mas is activated by ACE2 for an anti-inflammatory response to the infection via the ACE2/Ang(1-7)/Mas cascade. Therefore, we performed RT-qPCR on the same organs from the individuals of the three procedures of chronic ethanol exposure. Our results show a main effect of Group for Tmprss2, with a significant upregulation in the treatment animals of the non-abstinence vapor experiment (F[1, 43]=6.315, p=.016) with no interaction (F<1); the post-hoc test did not reveal any organ-specific effect (Fig. 2B). In the sub-chronic ethanol IP exposure procedure as well as in the post-dependent vapor treatment procedure no difference in Tmprss2 gene expression between treated and untreated animals could be detected (Fig. 2A, C). The non-abstinence animals represent the only condition of our study still containing significant blood alcohol concentrations (100-200 mg/dl) in the organism at time of death. Hence, these results suggest that Tmprss2 is regulated by acute ethanol intoxication. To verify this, we exposed animals to a single acute ethanol challenge IP of either low (0.5 g/kg), moderate (1.5 g/kg) or high dose (3.0 g/kg) (Suppl. Fig. 4). We found Tmprss2 mRNA levels to be differentially affected depending on dose and organ, with the liver showing increased levels only at the highest dose and the lung showing increased levels both at the moderate and high dose (Suppl. Fig. 1, Suppl. Tab. 9) confirming that an acute one-time ethanol treatment can increase Tmprss2 expression.

**Fig. 2:**
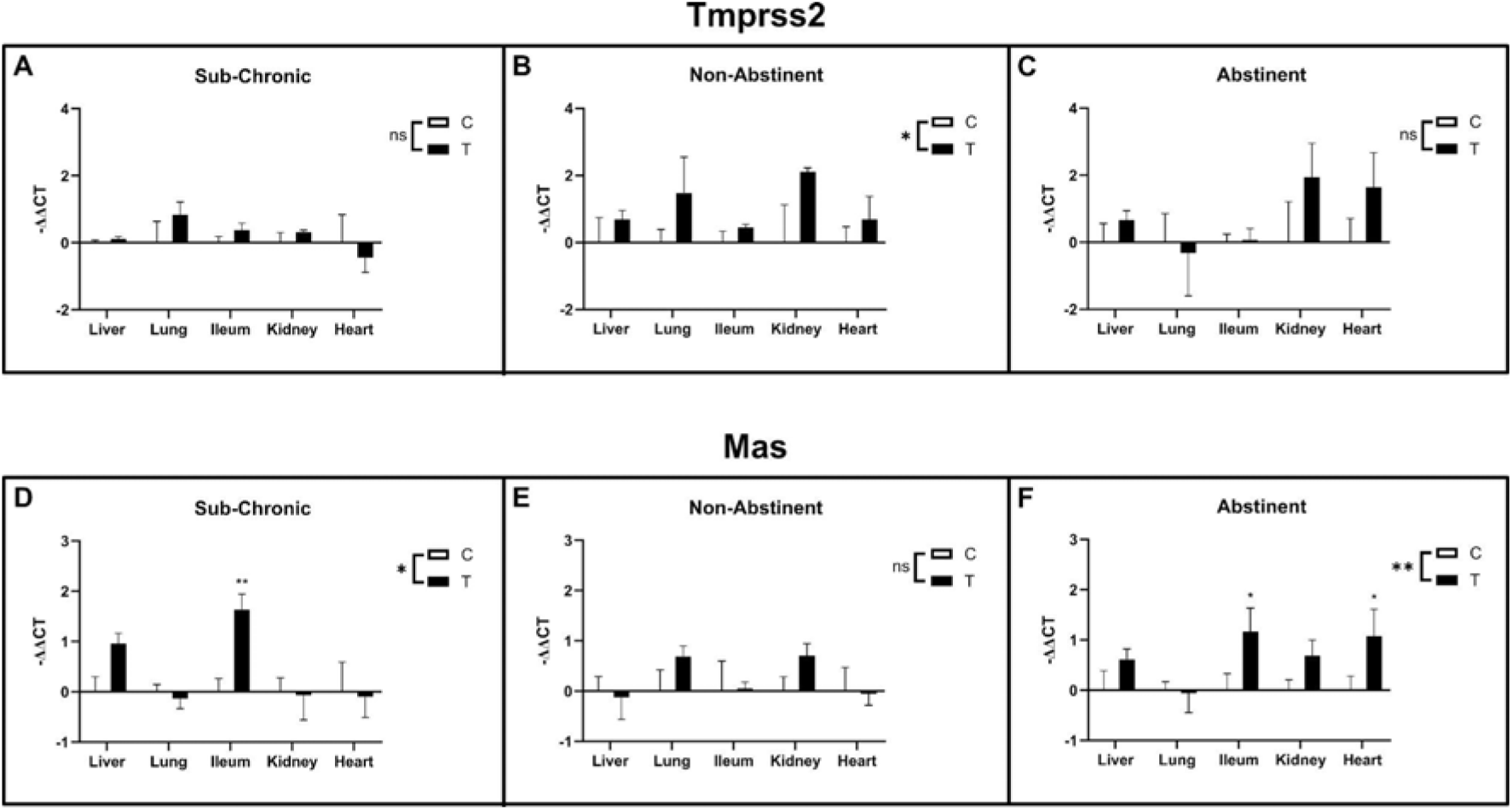
Gene expression of Tmprss2 (**A-C**) and Mas (**D-F**) in liver, lung, ileum, kidney and heart of sub-chronic ethanol IP injected and ethanol vapor exposed animals (indicated as “T”) with respective control groups (indicated as “C”). Data are shown as mean ± SE. For Tmprss2, factorial ANOVA indicated a treatment specific up-regulation in the non-abstinent vapor exposed group (**B**), whereas the sub-chronic ethanol IP injected group as well as the post-dependent (abstinence) groups were unaffected (**A** and **C**). Post-hoc tests in the NonAbst experiment did not reveal an organ-specific gene regulation. Mas was also treatment-specifically up-regulated. The sub-chronic as well as the post-dependent (abstinence) groups showed a significant difference compared to control in the factorial ANOVA (**D** and **F**). Subsequent post-hoc tests identified ileum as significantly up-regulated in either experiment. In addition, heart was up-regulated in the post-dependent (abstinence) condition (**F**).

For Mas gene expression, significant overall differences were identified in the sub-chronic treatment procedure (main effect of group: F[1, 49]=4.319, p=.043) (Fig. 2D) and the abstinence/post-dependent animals (main effect of group: F[1, 49]=9.778, p=.003) (Fig. 2F) with up-regulated Mas mRNA expression compared to their respective controls. In the subchronic treatment we also found a significant interaction (main effect of group: F[4, 49]=2.672, p=0.043), not present in the abstinence condition, indicating an organ-specific effect. Post-hoc analysis revealed that in the sub-chronic treatment condition Mas was significantly upregulated in the ileum (p=.002, d=2.32), while in the post-dependent (abstinence) condition Mas was upregulated in both the ileum (p=.022, d=1.17) and heart (p=.034, d=1.03). Since both treatments include an abstinence phase before sacrifice, the detected effects in gene expression might be related to the status of ethanol absence after long-term exposure. No effect of treatment or interaction was found in the non-abstinence condition (Fig. 2E). Thus, an augmented anti-inflammatory response to the infection via the ACE2/Ang(1-7)/Mas cascade might be the result of alcohol abstinence.

### Brain tissue seems to be less vulnerable to alcohol-induced changes of SARS-CoV2 infection-relevant genes

Several studies on patients that suffered from SARS-CoV2 infection reported neurological dysfunction as long-term side effects of the disease, even months after successful therapy (Ellul et al., 2020; Liu et al., 2021a; Mao et al., 2020; Moriguchi et al., 2020; Pajo et al., 2021; Paniz-Mondolfi et al., 2020). Furthermore, one of the first and most prominent symptoms of COVID-19 infection is anosmia (Hornuss et al., 2020). Therefore, the brain might be vulnerable to SARS-CoV2 infection as well. Given that alcohol induces long-lasting pronounced molecular changes in the brain following intermittent ethanol vapor exposure (Meinhardt et al., 2013, 2015; 2021b), we analyzed gene expression levels of Ace2, Tmprss2, and Mas in the olfactory bulb and the prefrontal cortex of ethanol vapor exposed animals (Fig. 3; Suppl. Tab. 4-6). As a second step, we performed in-situ gene expression analysis of the post-dependent animals to screen for further brain regions that might show differential expression due to long-term ethanol exposure (Fig. 3; Suppl. Tab. 7, 8). In the non-abstinence animals, Mas was downregulated specifically in the olfactory bulb, as indicated by the significant interaction (F[1, 16]=8.482, p=.010) and subsequent post-hoc (p=.006, d=1.68), while in the post-dependent group there was no difference between exposed and control animals (Fig. 3) suggesting that alcohol exposure briefly after a SARS-CoV2 infection may reduce an anti-inflammatory response via the ACE2/Ang(1-7)/Mas cascade – a mechanism that may contribute to the development of anosmia. Mas was unaffected in the abstinence condition (Fig. 3E). Ace2 and Tmprss2 mRNA levels were unaffected regardless of ethanol treatment conditions (Fig. 3A-D). This was further confirmed and extended by studying various brain regions via in-situ hybridization in the abstinence condition (post-dependent model). Thus, for Tmprss2 as well as Mas none of the 18 studied brain regions showed changes of transcript levels compared to the control group (Fig. 3 G-H; Suppl. Table 1), while Ace2 mRNA levels were below the detection limit; expression patterns are shown in Suppl. Fig. 3. In sum, we suggest that – except for the olfactory bulb - brain tissue seems to be less vulnerable to alcohol-induced changes of SARS-CoV2 infection-relevant genes.

**Fig. 3:**
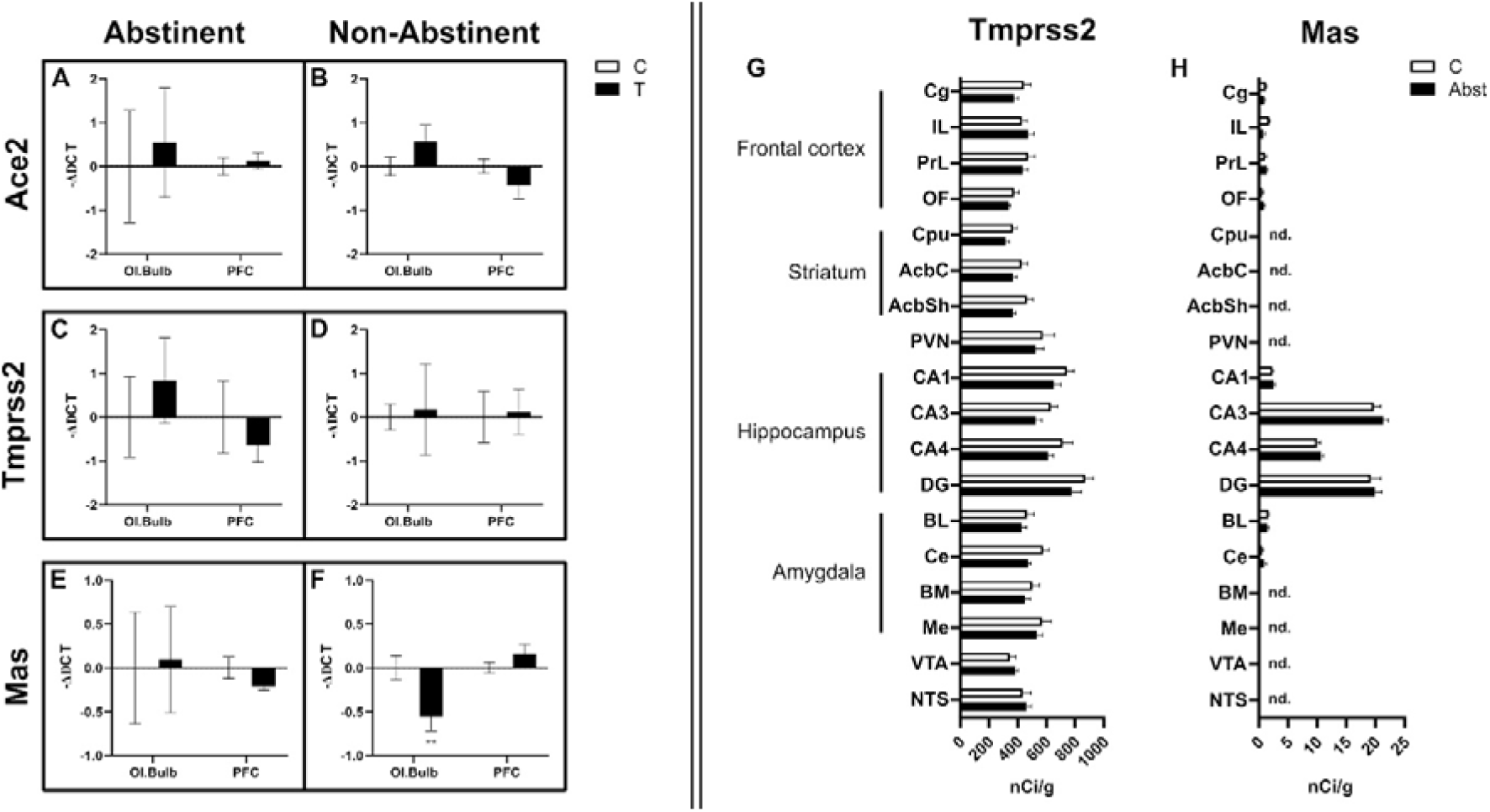
**A** Gene expression of Ace2 (**A-B**), Tmprss2 (**C-D**), and Mas (**E-F**) in the olfactory bulb and prefrontal cortex of ethanol vapor exposed animals measured via RT-qPCR. In the non-abstinent animals, solely the olfactory bulb showed significant down-regulation of Mas, whereas the remaining genes stayed unaffected. Transcript levels measured via in situ hybridization for Tmprss2 (**G**) and Mas (**H**) in 18 different brain regions of the ethanol vapor exposed with post-dependent (abstinence) condition relative to controls; no significant differential gene expression was detected in the studied brain regions. Ace2 signal was below the detection limit for all brain regions. From **A** to **F**, “T” indicates the treated group for each experiment (abstinent or non-abstinent), “C” indicates the respective control group. Data are shown as mean ± SE.

In summary, Mas gene expression was down-regulated in the olfactory bulb of vapor exposed rats (non-abstinence condition). In our brain-wide screening, we did not identify any other brain region with differentially regulated gene expression of SARS-CoV2 infection relevant target genes, suggesting the olfactory bulb as the main important brain region for SAR-CoV-2 infection.

## Discussion

In the present study we generated expression data of SARS-CoV2 infection-relevant genes (Ace2, Tmprss2 and Mas) in different organs following chronic alcohol exposure. We obtained three main findings. (i) Our results suggest an organ-specific differential regulation in a gene-specific manner, where Ace2 is consistently up-regulated in the lung across all three test conditions. This finding suggests a higher abundance of ACE2 protein in lung tissue following chronic alcohol intake and therefore an increased stochastic probability of the SARS-CoV2 to enter and infect the cells. Hence, chronic alcohol consumption is a risk factor for COVID-19, at least on the molecular level. (ii) Following alcohol abstinence, we observed an augmented anti-inflammatory response via the ACE2/Ang(1-7)/Mas cascade, which suggests that after abstinence a protective mechanism may kick in. However, (iii) in the olfactory bulb an opposing effect was observed. There was a down-regulation of Mas gene expression that might lead to a lower activation of the anti-inflammatory ACE2/Ang(1-7)/Mas cascade after viral infection, which might intensify the disease progression towards anosmia.

Previous studies have already shown that patients with substance use disorder (SUDs) are at a higher risk of developing COVID-19 and have a worse prognosis including death (Baillargeon et al., 2021; Patanavanich & Glantz, 2021; Wang et al., 2021). Up-regulation of Ace2 has also been shown in SUD patients (Althobaiti et al., 2021; Dubey et al., 2020; Mallet et al., 2020; Wei & Shah, 2020). Furthermore, increased Ace2 expression levels have been reported in cigarette smokers (Smith et al., 2020). Our study expands on these findings, indicating that a similar effect might be also the case for chronic alcohol consumption. Moreover, our results is in line with previous findings of increased Ace2 expression in the brain of AUD patients (Muhammad et al., 2021) and suggest that such an effect may not only be confined to the brain. Additionally, it is well known that most of the AUD patients also smoke regularly (Bobo & Husten, 2000; Grant et al., 2004), which has been shown to have a negative impact on the immune system and increases the pathogenic infection risk in general (Qiu et al., 2017).

The lung is the most vulnerable organ for COVID-19 infection, as it represents the main route of the infection (Jiang et al., 2020; D. Wang et al., 2020; Yang et al., 2020). Up to date, it is not completely clear which influence increased ACE2 expression levels have in the context of SARS-CoV2 infection and COVID-19 prognosis (Ni et al., 2020). However, SARS-CoV - a closely related infectious virus - more readily infects lung cells with higher Ace2 expression levels (Jia et al., 2005). It is plausible to think that such a phenomenon could also occur for SARS-CoV2. Furthermore, it was shown that elevated Ace2 expression levels in different human organs correlate with higher infection probability of those same organs (He et al., 2021). SARS-CoV2 infection, once occurred, has been shown to induce Ace2 downregulation by internalization during virus entry. This in turn induces an imbalance in the ACE2/Ang(1-7)/Mas axis determining an overactivation of the ACE/AngII/AT1R axis leading finally to inflammatory response, oxidative stress and ultimately to ARDS and, in the worst cases, systemic organ failure and death (Kaseb et al., 2021). A recent meta-analysis focusing on the expression profiles of Ace2 in several human tissues found that Ace2, as Tmprss2, appear to be induced by pro-inflammatory states found in conditions such as obesity and diabetes (Gkogkou et al., 2020), which also represent risk factors for SARS-CoV2 infection and severe COVID-19 (Hamer et al., 2020; Zheng et al., 2020).

Ace2 upregulation as a consequence of chronic alcohol intake could be one potential explanation for epidemiological data revealing increased infection rates among AUD patients and worsened COVID-19 prognosis. The amount of alcohol consumed per week was in fact found to be correlated to ARDS after SARS-CoV2 infection in a cohort study (Lassen et al., 2021). Furthermore, studies have found associations between AUD and hospitalization risk after SARS-CoV2 infection (Allen et al., 2020), risk of prolonged hospital stay and need of respiratory support (Varela Rodriguez et al., 2021). Chronic alcohol consumption is also known to induce disruption of the innate and adaptive immune system (Boé et al., 2009; Downs et al., 2013; Kershaw & Guidot, 2008; Simou et al., 2018), which represent a further factor that could cause a predisposition for severe SARS-CoV2 infection. However, it should be noted that a correlation between alcohol consumption and risk of severe SARS-CoV2 infection was not found in a cohort study based on over 300.000 adults observing COVID-19-related hospitalization rate in the UK (Hamer et al., 2020).

ACE2 is not the only important component for SARS-CoV2 viral infection. TMPRSS2 is also involved in facilitating virus entry by cleaving its spike proteins that are recognized and bound by ACE2 (Hoffmann et al., 2020). There was an overall upregulation of Tmprss2 in the non-abstinence ethanol vapor exposed rats without any organ-specific effects. Therefore, based on our results, Tmprss2 mRNA expression seems to be influenced by relevant blood alcohol concentrations – an assumption that is supported by our additional experiments where alcohol was given acutely. Since TMPRSS2 is supporting SARS-CoV2 cell entry, its upregulation might accelerate the process of virus infiltration. Hence, TMPRSS2 inhibitors are discussed as promising therapeutic approaches for SARS-CoV2 infection (Hoffmann et al., 2020; McKee et al., 2020; Vahed et al., 2020).

The G-protein coupled receptor Mas is also a key player in the SARS-CoV2 infection pathway, as part of the ACE2/Ang1-7/Mas cascade (Gheblawi et al., 2020). Once ACE2 has catalyzed the reaction of Ang II to Ang1-7 on the cell surface, the cell-membrane bound receptor Mas activates an anti-inflammatory cascade inside the cell. In our study, Mas showed treatment-specific upregulation in the ileum of animals that underwent sub-chronic ethanol IP treatment as well as ethanol vapor exposure– in both groups, rats underwent abstinence before organ removal. The ileum is known to be vulnerable to SARS-CoV2 infection. Infected patients report diarrhea, nausea and vomiting mainly in the beginning of disease progression and the gastrointestinal tract is suggested as an alternative pathway of SARS-CoV2 infection (Gu et al., 2020; Kotfis & Skonieczna-Żydecka, 2020). The Mas receptor represents the counter regulator part of the RAS including vasodilatation, antiproliferation, anti-fibrosis, cell proliferation, and fluid volume homeostasis (Vaajanen et al., 2015). In this context, an upregulation of Mas might lead to a faster and more efficient direct cell response to the infection and therefore could contribute to a protective effect for severe COVID-19 (Gheblawi et al., 2020).

In the brain, we observed down-regulation of Mas in the olfactory bulb of ethanol vapor exposed rats (post-dependent model). Down-regulation of Mas gene expression might lead to a lower activation of the anti-inflammatory ACE2/Ang(1-7)/Mas cascade after viral infection, which in turn may result in anosmia. Beside the Mas alteration in the olfactory bulb, brain tissue in general seems to be less vulnerable to alcohol-induced changes of SARS-CoV2 infection-relevant genes. A finding which is in line with the low overall Ace2 expression levels in the rodent brain – with the exception of the olfactory bulb that shows the highest Ace2 gene expression across the rodent brain (https://mouse.brain-map.org/gene/show/45849). Thus, chronic alcohol consumption may constitute a risk factor for the development of anosmia during a SARS-CoV2 infection. This conclusion is supported by previous studies that showed abnormal development of the olfactory bulb in animals exposed to alcohol prenatally or in neonatal stage (Akers et al., 2011; Burke et al., 2016; Chen et al., 1999). In adult animals, the functionality of the olfactory bulb is also known to be impaired by alcohol consumption (Collins et al., 1998; Gericke et al., 2006; Penland et al., 2001). It would be interesting to examine the occurrence of anosmia in large-scale cohorts of Covid-19 patients in relation to alcohol consumption. In this respect, it is of note that anosmia was not recognized as a prominent sign of COVID-19 infection in infected Chinese from Wuhan where usually alcohol drinking is much less common than in European or US population (Izquierdo-Dominguez et al., 2020; Mullol et al., 2020).

In conclusion, our study provides evidence on the molecular level that chronic alcohol consumption constitutes a potential risk factor for SARS-CoV2 infection. However, this study represents only an indirect indication of potential vulnerability to SARS-CoV2 infection in individuals who chronically consume alcohol and does not allow any conclusion on severity of Covid-19 disease progression - with one exception that chronic alcohol consumption may promote anosmia via downregulation of the ACE2/Ang(1-7)/Mas cascade after viral infection. To fully state the potential impact of chronic alcohol consumption on SARS-CoV2 infection risk and Covid-19 disease outcome, additional research needs to be performed. We suggest analysis of SARS-CoV2 infection-relevant genes in alcohol-exposed rodents which are infected with SARS-CoV2.

## Materials and Methods

### Sub-chronic ethanol IP treatment

Seven weeks old male Wistar rats (Envigo, Ettlingen, Germany) were randomly distributed in groups and cages of three animals per cage. The animals were habituated to the housing conditions for one week. Afterwards, two weeks of daily ethanol injections (1.5 g/kg; intraperitoneal (IP)) were applied twice a day. The first injection was 2 h before the beginning of the active cycle of the animals, the second injection was given 6 h after the first. In total, 6 animals were used for the ethanol IP treatment, and additional six control animals received repetitive saline injections. The animals were sacrificed 6 h after the last IP injection via decapitation, during the active cycle (ZT22), since previous studies have shown that the circadian regulation of Ace2 peaks at this time point (Herichová et al., 2013; Zlacká et al., 2021). The organs lung, liver, ileum, kidney, and heart were dissected right after, snap frozen in isopentane and stored in −80° C until further processing. These organs were chosen, because SARS-CoV2 genome has been detected in all of them in human patients (Li et al., 2020; Puelles et al., 2020; Xiao et al., 2020; Zou et al., 2020), showing that they represent potential sites for SARS-CoV2 infection and replication within the human body. Furthermore, all of these organs are known to be affected by chronic alcohol consumption in humans (Epstein, 1997; Osna et al., 2017; Piano, 2017; Simet & Sisson, 2015; Y. Wang et al., 2014). Figure 4 represents the overview of the experimental design. All experimental procedures were approved by Regierungspräsidium Karlsruhe, Germany (license: G-208/16).

**Fig. 4:**
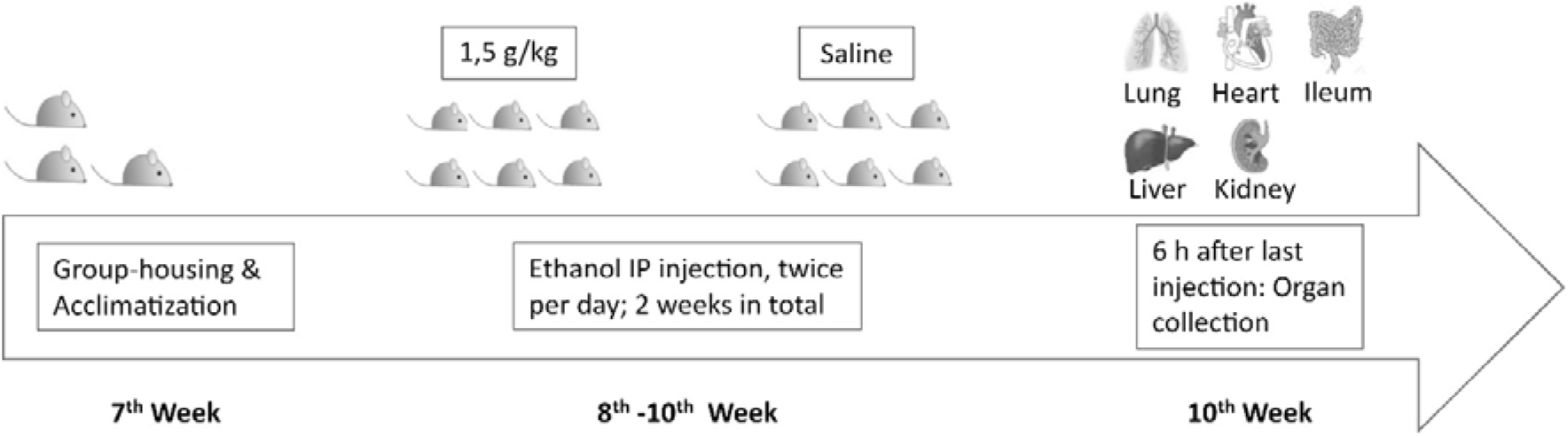
Time-table of sub-chronic ethanol IP injections in male Wistar rats. The animals arrived in the laboratory at the age of 7 weeks. After one week of acclimatization to the housing room, two weeks of daily ethanol IP injection were given. Lung, heart, ileum, liver, and kidney were dissected 6 h after the last injection.

### Chronic intermittent ethanol vapor exposure

The procedure of chronic intermittent ethanol exposure is a well-established animal model of alcohol dependence ((Meinhardt & Sommer, 2014); license: G-10/16). In our experiment, male Wistar rats (Envigo, Ettlingen, Germany) at the age of seven weeks were habituated to the experimental conditions for one week. Then, the animals were randomly distributed into treatment and control groups. The treatment group was exposed to ethanol vaporized air in a vapor chamber for seven consecutive weeks for 16 h per day. Control animals were housed in the same room, but were exposed to room air only. Twice a week, blood alcohol concentrations (BAC) were measured directly after the exposure cycle using approximately 10 μl blood of the animals’ tail tip (values see Suppl. Fig 2). Vapor exposure was adjusted to the measured BAC values to keep them between 150 and 300 mg/dl during the whole exposure period. Seven hours after the end of the penultimate vapor exposure, withdrawal scores were measured, based on a previously published method (Macey et al., 1996). Two hours before the end of the last ethanol exposure half of the exposed and half of the control animals were sacrificed, representing the acute effect of chronic ethanol exposure (non-abstinence group). After three weeks of abstinence, the remaining half of the batch was sacrificed, which represents the post-dependent state of chronic ethanol exposure (Meinhardt & Sommer, 2014) (abstinence group). All animals were sacrificed at the same time point as the ethanol IP injected animals: 2 h before the end of the active cycle (lights on: 3 am, lights off: 3 pm; ZT22; Fig. 5). During the alcohol vapor exposure procedure, one animal died resulting in a final sample size of n = 5 + 6 rats for treated and controls respectively in the “non-abstinence” condition, and n = 6 + 6 rats for treated and controls in the abstinence condition. Details of the experimental setup can be found in Fig. 5.

**Fig. 5:**
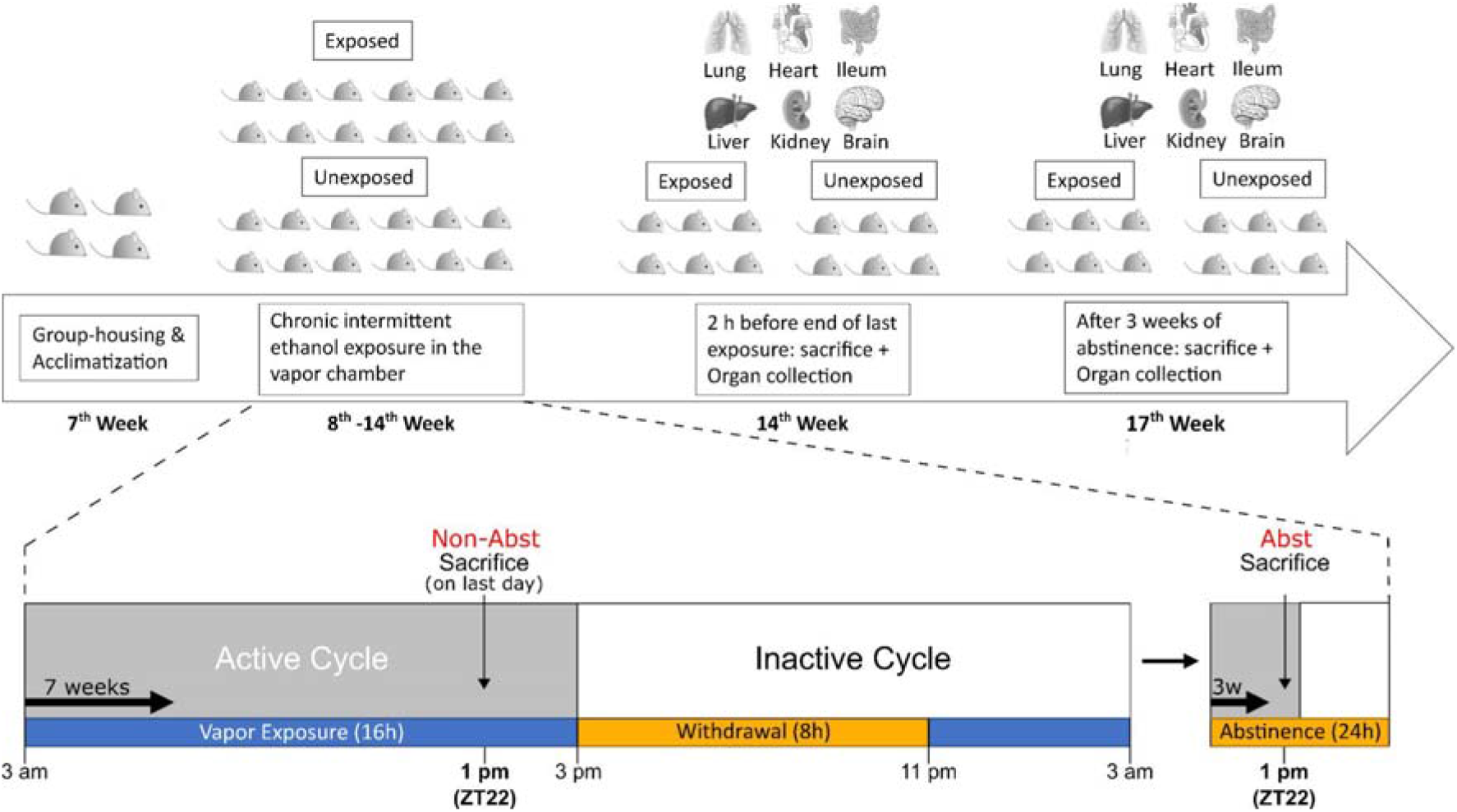
Timeline of the chronic intermittent ethanol vapor exposure experiment. **Top**: General experimental setup. Daily cycles of ethanol vapor exposure were performed for seven consecutive weeks with 16 h exposure per day. During the last exposure, half of the animals were sacrificed and the respective organs were collected. After three weeks of abstinence, the remaining half of the batch was sacrificed, including organ collection. **Bottom**: exposure and recovery times during the seven weeks of intermittent vapor exposure. The animals were sacrificed two hours before end of the active cycle (ZT22), matching with the sub-chronic ethanol IP treated animals.

### RNA extraction and qRT-PCR

Total RNA of kidney, lung, heart, liver, ileum and brain of the sub-chronic IP injected animals as well as the CIE rats was extracted using Trizol (Thermo Fisher, Waltham, MA, USA) and the RNeasy MicroElute RNA extraction kit (Qiagen, Hilden, Germany). After reverse transcription into cDNA using High-Capacity cDNA Reverse Transcription Kit (Thermo Fisher, Waltham, MA, USA), the samples were randomized on RT-qPCR plates and analyzed for gene expression of Ace2, Tmprss2, and Mas relative to one housekeeping gene (Gapdh). The sequences of the RT-qPCR primer pairs are listed in Suppl. Tab. 10. All samples were pipetted in three technical replicates. Technical outlier was defined as deviation of greater than 5% from the mean of the triplicates.

### In-situ hybridization in rat brain tissue

Coronal brain slices (10 μm thick) from male Wistar rats after ethanol vapor exposure were sliced using a Cryostat. We obtained samples from 18 different brain regions: cingulate cortex (Cg), infralimbic cortex (IL), prelimbic cortex (PrL), orbitofrontal cortex (OF), caudate putamen (CPu), nucleus accumbens core (AcbC) and shell (AcbSh), paraventricular nucleus (PVN), cornu ammonis area 1 (CA1), cornu ammonis area 3 (CA3), cornu ammonis area 4 (CA4), dentate gyrus (DG), basolateral amygdala (BLA), central amygdala (CeA), basomedial amygdala (BMA), medial amygdala (MeA), ventral tegmental area (VTA), nucleus tractus solitarii (NTS). The detailed procedure is described in the Supplementary section. Reference for comparison of the expression patterns was taken from The Allen Brain Atlas and cited studies from the website (https://mouse.brain-map.org/gene/show/30018).

### Statistical analysis

RT-qPCR data were analyzed separately for each gene of interest (Ace2, Tmprss2, Mas). Firstly, Gapdh Ct values were analyzed to make sure that no group difference was present. dCt values for each gene of interest were then obtained and extreme outliers (greater or lower than three times the interquartile range) were excluded. For organ-wise analysis, dCt values were then used to perform a Factorial (Two-Way) ANOVA using group (control or treated) and organ (liver, lung, ileum, kidney, heart) as the two categorical factors. Olfactory bulb and PFC data were analyzed via Factorial (Two-Way) ANOVA using group (control or treated) and region (Olf. bulb, PFC) as the two categorical factors. Depending on the experiment, the treated group was either sub-chronic, non-abstinence, or abstinence. Fisher LSD was used for post-hoc analysis. Due to the exploratory nature of our study, we conducted the post-hoc test also when a significant main effect of the group was detected without a significant interaction, in order to increase the information gain from the experiments. Alongside the p-value, we also report the effect size (Cohen’s d) for every organ or brain region found to be significantly different from the control condition according to the post-hoc test.

In situ data (measured in nCi/g) from each brain region were analyzed via multiple t-tests. Nanocurie per gram (nCi/g) values were obtained after conversion of the final mean gray values measured with MCID Analysis 7.1. Technical outliers were identified as greater or lower than two times the standard deviation. Biological outliers were defined as greater or lower than three times the interquartile range.

Statistica10 (StatSoft) and GraphPad Prism 8.4.3 were used for statistical analysis and graphs.

## Acknowledgement

Thanks for Rick Bernardi for professional English editing.

## Author contribution

MMF, FG, ACH and RS designed the experiments. MMF, FG, MS and RoS performed experiments. MMF, FG and MS analyzed the data. MMF and RS wrote the manuscript. RS and ACH provided financial and scientific support.

