## Supplementary for "Chronic alcohol intake regulates expression of SARS-CoV2 infection-relevant genes in an organ-specific manner"

**Supplementary results**

**Acute, one-time ethanol IP treatment**

Tmprss2 mRNA expression levels measured via RT-qPCR in liver, lung, and ileum after a single ethanol IP injection at different doses (0.5 g/kg, 1.5 g/kg, 3.0 g/kg; Suppl. Fig. 3) were analyzed via Factorial (Two-Way) ANOVA. We found a significant interaction between dose and organ (F[6, 60]=2.444, p=.035), as well as a significant main effect of dose (F[3, 60]=3.107, p=.033) and organ (F[2, 60]=1015.1, p=.000). After post-hoc analysis of the interaction, we found Tmprss2 mRNA levels to be significantly increased in the liver at the highest dose (p=.0502, d=1.67). Tmprss2 mRNA was instead up-regulated after both at 1.5 g/kg (p=.000, d=1.57) and 3.0 g/kg (p=.046, d=0.69) single injection. Tmprss2 was not significantly affected in the ileum (Suppl. Fig. 1; Suppl. Tab. 6).

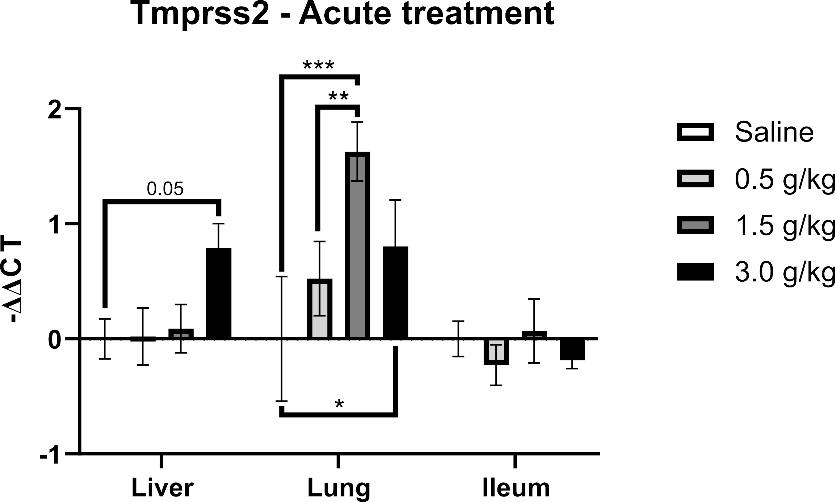

**Suppl. Fig. 1**: Tmprss2 gene expression in liver, lung, and ileum after a one-time ethanol IP injection at three different doses (0.5 g/kg, 1.5 g/kg, 3.0 g/kg). Each treatment group is represented by 6 animals. Data are shown as mean ± SE.

**Ethanol vapor treatment with BAC and withdrawal measurement**

The Blood alcohol concentration (BAC) was measured twice a week to adjust the pump rate to maintain the BAC of the animals between 150 and 300 mg/dl (Suppl. Fig. 2A). Withdrawal signs scored were ventral limb retraction (VLR), vocalization, tail stiffness, tremors and abnormal gait (Suppl. Fig. 2B). After adding the different withdrawal signs for each animal (cumulative withdrawal score) we found that alcohol exposed animals (mean= 3.72, median = 4) showed greater signs of withdrawal when compared to the air exposed control group (mean = 1.83, median = 1.5), as indicated by Mann-Whitney U test (U = 22.50, z = -2.65, p = .008, r = -.55; Suppl. Fig. 2C).

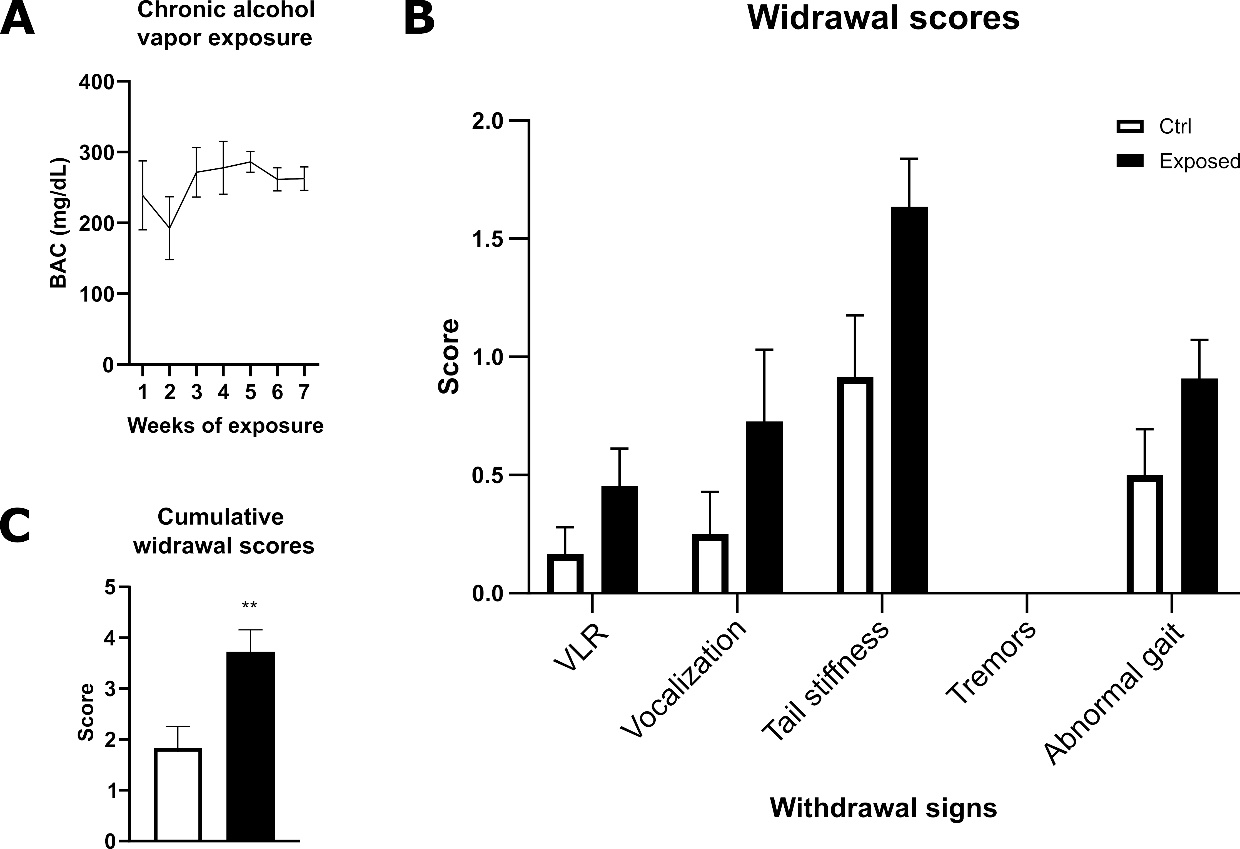

**Suppl. Fig. 2**: Blood alcohol concentration (BAC) throughout the seven weeks of alcohol vapor exposure with 16 h exposure period per day (**A**). BAC was measured twice per week by puncturing the tip of the tail from three random animals each time, in order to adjust the pump rate to maintain the BAC between 150 and 300 mg/dL Withdrawal signs scores for the alcohol exposed and control animals (**B**), the scored signs were ventral limbic retraction (VLR), vocalization, tail stiffness, tremors, and abnormal gait. Seven hours after the end of the second last day of exposure the withdrawal signs were scored for all animals (the experimenter who performed the scoring was blinded to the groups). Cumulative withdrawal scores for alcohol exposed and control animals (**C**). Results are shown as mean ± standard error.

**Tmprss2 and Mas expression pattern**

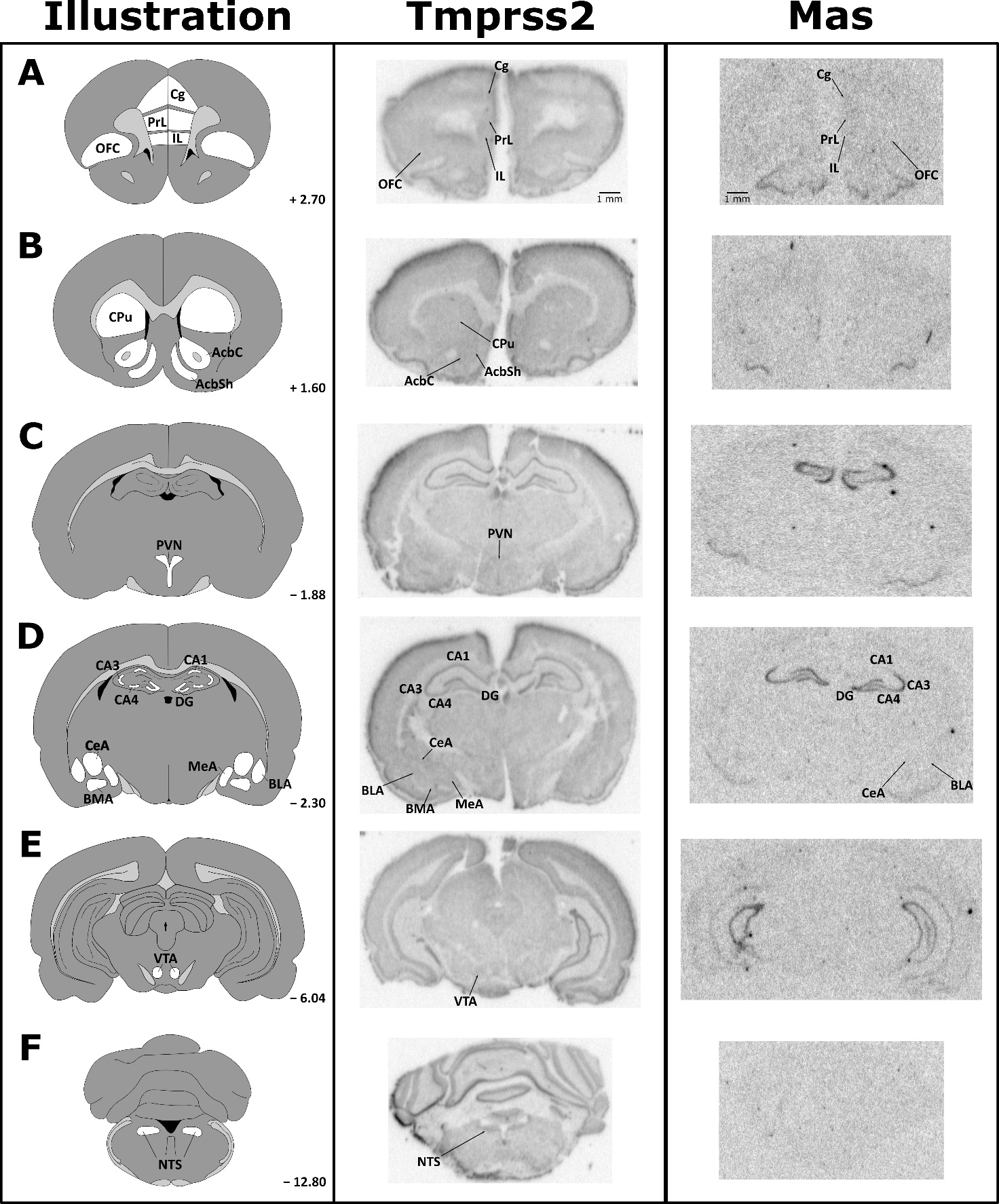

**Suppl. Fig. 3:** Schematic illustration (left) of coronal rat brain sections with the brain regions analyzed via in situ hybridization according to Paxinos and Watson (1998). Representative autoradiograms showing Tmprss2 (center) and Mas (right) mRNA expression patterns. The regions analyzed were the prefrontal cortex (A): specifically cingulate cortex (Cg), prelimbic cortex (PrL), infralimbic cortex (IL), orbitofrontal cortex (OFC); the striatum (B): specifically caudate putamen (CPu), nucleus accumbens core (AcbC) and shell (AcbSh); the paraventricular nucleus (PVN) (C); the hippocampus and the amygdala (D): specifically cornu ammonis area 1 (CA1), cornu ammonis area 3 (CA3), cornu ammonis area 4 (CA4), dentate gyrus (DG), basolateral amygdala (BLA), central amygdala (CeA), basomedial amygdala (BMA), medial amygdala (MeA); the ventral tegmental area (VTA) (E), and the nucleus tractus solitarii (NTS) (F).

**Additional analyzes of the RT-qPCR data of Ace2, Mas and Tmprss2 in peripheral organs of sub-chronic ethanol IP treatment and ethanol vapor exposed animals**

**Suppl. Tab. 1**: Factorial ANOVA of Ace2 dCt values in sub-chronic, non abstinent, and abstinent models

| Sub-chronic ethanol IP treatment | | | | | | | | | |
| --- | --- | --- | --- | --- | --- | --- | --- | --- | --- |
| Descriptives | | | | Statistics | | | | | |
| **Organ** | **Group** | **dCt ± SE** | **n** | **Predictor** | **Degrees of Freedom** | **Mean Square** | **F-value** | **p-value** | **Partial η2** |
| Liver | Ctrl | 11.52 ± 0.69 | 6 | Organ | 4 | 360.042 | 299.894 | 0.0000 | 0.9615 |
|  | SubChr | 9.63 ± 0.05 | 6 | Group | 1 | 6.191 | 5.157 | 0.0277 | 0.0970 |
| Lung | Ctrl | 6.19 ± 0.46 | 6 | Organ*Group | 4 | 3.680 | 3.065 | 0.0250 | 0.2035 |
|  | SubChr | 4.29 ± 0.09 | 5 | Error | 48 | 1.201 |  |  |  |
| Ileum | Ctrl | -0.41 ± 0.28 | 6 |  |  |  |  |  |  |
|  | SubChr | -0.36 ± 0.30 | 6 |  |  |  |  |  |  |
| Kidney | Ctrl | 7.72 ± 0.47 | 6 |  |  |  |  |  |  |
|  | SubChr | 7.98 ± 0.48 | 6 |  |  |  |  |  |  |
| Heart | Ctrl | 14.33 ± 0.77 | 6 |  |  |  |  |  |  |
|  | SubChr | 14.53 ± 0.07 | 5 |  |  |  |  |  |  |
| Chronic ethanol vapor without abstinence | | | | | | | | | |
| Descriptives | | | | Statistics | | | | | |
| **Organ** | **Group** | **dCt ± SE** | **n** | **Predictor** | **Degrees of Freedom** | **Mean Square** | **F-value** | **p-value** | **Partial η2** |
| Liver | Ctrl | 11.93 ± 0.71 | 6 | Organ | 4 | 397.944 | 99.490 | 0.0000 | 0.9025 |
|  | NonAbst | 10.81 ± 0.14 | 4 | Group | 1 | 29.759 | 7.440 | 0.0092 | 0.1475 |
| Lung | Ctrl | 9.70 ± 0.18 | 5 | Organ*Group | 4 | 2.870 | 0.718 | 0.5846 | 0.0626 |
|  | NonAbst | 6.68 ± 1.39 | 5 | Error | 43 | 4.000 |  |  |  |
| Ileum | Ctrl | 0.82 ± 0.48 | 6 |  |  |  |  |  |  |
|  | NonAbst | -0.57 ± 0.41 | 5 |  |  |  |  |  |  |
| Kidney | Ctrl | 15.18 ± 1.40 | 6 |  |  |  |  |  |  |
|  | NonAbst | 13.30 ± 1.38 | 5 |  |  |  |  |  |  |
| Heart | Ctrl | 15.18 ± 0.35 | 6 |  |  |  |  |  |  |
|  | NonAbst | 15.03 ± 0.58 | 5 |  |  |  |  |  |  |
| Chronic ethanol vapor with abstinence | | | | | | | | | |
| Descriptives | | | | Statistics | | | | | |
| **Organ** | **Group** | **dCt ± SE** | **n** | **Predictor** | **Degrees of Freedom** | **Mean Square** | **F-value** | **p-value** | **Partial η2** |
| Liver | Ctrl | 12.80 ± 1.08 | 6 | Organ | 4 | 484.672 | 120.498 | 0.0000 | 0.9077 |
|  | Abst | 12.13 ± 0.86 | 6 | Group | 1 | 31.882 | 7.926 | 0.0070 | 0.1392 |
| Lung | Ctrl | 7.42 ± 1.18 | 6 | Organ*Group | 4 | 8.525 | 2.120 | 0.0925 | 0.1475 |
|  | Abst | 5.05 ± 0.38 | 6 | Error | 49 | 4.022 |  |  |  |
| Ileum | Ctrl | -0.48 ± 0.57 | 6 |  |  |  |  |  |  |
|  | Abst | -0.33 ± 0.53 | 6 |  |  |  |  |  |  |
| Kidney | Ctrl | 15.25 ± 1.51 | 6 |  |  |  |  |  |  |
|  | Abst | 11.18 ± 0.39 | 5 |  |  |  |  |  |  |
| Heart | Ctrl | 15.34 ± 0.21 | 6 |  |  |  |  |  |  |
|  | Abst | 14.93 ± 0.31 | 6 |  |  |  |  |  |  |

**Suppl. Tab. 2**: Factorial ANOVA of Mas dCt values in sub-chronic, non abstinent, and abstinent models

| Sub-chronic ethanol IP treatment | | | | | | | | | |
| --- | --- | --- | --- | --- | --- | --- | --- | --- | --- |
| Descriptives | | | | Statistics | | | | | |
| **Organ** | **Group** | **dCt ± SE** | **n** | **Predictor** | **Degrees of Freedom** | **Mean Square** | **F-value** | **p-value** | **Partial η2** |
| Liver | Ctrl | 14.39 ± 0.30 | 5 | Organ | 4 | 196.703 | 274.728 | 0.0000 | 0.9573 |
|  | SubChr | 13.43 ± 0.21 | 6 | Group | 1 | 3.092 | 4.319 | 0.0430 | 0.0810 |
| Lung | Ctrl | 8.79 ± 0.15 | 6 | Organ*Group | 4 | 1.913 | 2.672 | 0.0429 | 0.1790 |
|  | SubChr | 8.93 ± 0.19 | 6 | Error | 49 | 0.716 |  |  |  |
| Ileum | Ctrl | 14.32 ± 0.27 | 6 |  |  |  |  |  |  |
|  | SubChr | 12.68 ± 0.31 | 6 |  |  |  |  |  |  |
| Kidney | Ctrl | 5.46 ± 0.28 | 6 |  |  |  |  |  |  |
|  | SubChr | 5.53 ± 0.49 | 6 |  |  |  |  |  |  |
| Heart | Ctrl | 5.51 ± 0.59 | 6 |  |  |  |  |  |  |
|  | SubChr | 5.61 ± 0.41 | 6 |  |  |  |  |  |  |
| Chronic ethanol vapor without abstinence | | | | | | | | | |
| Descriptives | | | | Statistics | | | | | |
| **Organ** | **Group** | **dCt ± SE** | **n** | **Predictor** | **Degrees of Freedom** | **Mean Square** | **F-value** | **p-value** | **Partial η2** |
| Liver | Ctrl | 13.57 ± 0.29 | 6 | Organ | 4 | 248.273 | 314.813 | 0.0000 | 0.9662 |
|  | NonAbst | 13.69 ± 0.44 | 5 | Group | 1 | 0.857 | 1.087 | 0.3028 | 0.0241 |
| Lung | Ctrl | 2.60 ± 0.42 | 6 | Organ*Group | 4 | 0.451 | 0.572 | 0.6846 | 0.0494 |
|  | NonAbst | 1.91 ± 0.21 | 5 | Error | 44 | 0.789 |  |  |  |
| Ileum | Ctrl | 11.87 ± 0.60 | 6 |  |  |  |  |  |  |
|  | NonAbst | 11.81 ± 0.13 | 5 |  |  |  |  |  |  |
| Kidney | Ctrl | 5.79 ± 0.29 | 6 |  |  |  |  |  |  |
|  | NonAbst | 5.08 ± 0.24 | 5 |  |  |  |  |  |  |
| Heart | Ctrl | 5.54 ± 0.47 | 6 |  |  |  |  |  |  |
|  | NonAbst | 5.60 ± 0.23 | 4 |  |  |  |  |  |  |
| Chronic ethanol vapor with abstinence | | | | | | | | | |
| Descriptives | | | | Statistics | | | | | |
| **Organ** | **Group** | **dCt ± SE** | **n** | **Predictor** | **Degrees of Freedom** | **Mean Square** | **F-value** | **p-value** | **Partial η2** |
| Liver | Ctrl | 14.43 ± 0.39 | 6 | Organ | 4 | 300.465 | 410.605 | 0.0000 | 0.9710 |
|  | Abst | 13.81 ± 0.21 | 6 | Group | 1 | 7.155 | 9.778 | 0.0030 | 0.1664 |
| Lung | Ctrl | 2.08 ± 0.17 | 5 | Organ*Group | 4 | 0.667 | 0.911 | 0.4648 | 0.0692 |
|  | Abst | 2.14 ± 0.39 | 6 | Error | 49 | 0.732 |  |  |  |
| Ileum | Ctrl | 12.55 ± 0.33 | 6 |  |  |  |  |  |  |
|  | Abst | 11.38 ± 0.47 | 6 |  |  |  |  |  |  |
| Kidney | Ctrl | 5.86 ± 0.21 | 6 |  |  |  |  |  |  |
|  | Abst | 5.17 ± 0.31 | 6 |  |  |  |  |  |  |
| Heart | Ctrl | 5.47 ± 0.28 | 6 |  |  |  |  |  |  |
|  | Abst | 4.39 ± 0.53 | 6 |  |  |  |  |  |  |

**Suppl. Tab. 3**: Factorial ANOVA of Tmprss2 dCt values in sub-chronic, non abstinent, and abstinent models

| Sub-chronic ethanol IP treatment | | | | | | | | | |
| --- | --- | --- | --- | --- | --- | --- | --- | --- | --- |
| Descriptives | | | | Statistics | | | | | |
| **Organ** | **Group** | **dCt ± SE** | **n** | **Predictor** | **Degrees of Freedom** | **Mean Square** | **F-value** | **p-value** | **Partial η2** |
| Liver | Ctrl | 9.41 ± 0.08 | 5 | Organ | 4 | 327.950 | 312.659 | 0.0000 | 0.9638 |
|  | SubChr | 9.30 ± 0.08 | 5 | Group | 1 | 0.822 | 0.783 | 0.3810 | 0.0164 |
| Lung | Ctrl | 12.85 ± 0.64 | 6 | Organ*Group | 4 | 0.635 | 0.606 | 0.6604 | 0.0490 |
|  | SubChr | 12.01 ± 0.38 | 6 | Error | 47 | 1.049 |  |  |  |
| Ileum | Ctrl | 2.93 ± 0.19 | 6 |  |  |  |  |  |  |
|  | SubChr | 2.54 ± 0.21 | 6 |  |  |  |  |  |  |
| Kidney | Ctrl | 5.89 ± 0.31 | 6 |  |  |  |  |  |  |
|  | SubChr | 5.58 ± 0.07 | 5 |  |  |  |  |  |  |
| Heart | Ctrl | 15.78 ± 0.84 | 6 |  |  |  |  |  |  |
|  | SubChr | 16.22 ± 0.45 | 6 |  |  |  |  |  |  |
| Chronic ethanol vapor without abstinence | | | | | | | | | |
| Descriptives | | | | Statistics | | | | | |
| **Organ** | **Group** | **dCt ± SE** | **n** | **Predictor** | **Degrees of Freedom** | **Mean Square** | **F-value** | **p-value** | **Partial η2** |
| Liver | Ctrl | 9.48 ± 0.74 | 6 | Organ | 4 | 335.333 | 140.935 | 0.0000 | 0.9291 |
|  | NonAbst | 8.79 ± 0.26 | 5 | Group | 1 | 15.024 | 6.314 | 0.0158 | 0.1280 |
| Lung | Ctrl | 5.16 ± 0.39 | 6 | Organ*Group | 4 | 1.104 | 0.464 | 0.7618 | 0.0414 |
|  | NonAbst | 3.68 ± 1.07 | 5 | Error | 43 | 2.379 |  |  |  |
| Ileum | Ctrl | 2.98 ± 0.34 | 6 |  |  |  |  |  |  |
|  | NonAbst | 2.52 ± 0.09 | 5 |  |  |  |  |  |  |
| Kidney | Ctrl | 9.04 ± 1.12 | 6 |  |  |  |  |  |  |
|  | NonAbst | 6.93 ± 0.11 | 3 |  |  |  |  |  |  |
| Heart | Ctrl | 17.38 ± 0.47 | 6 |  |  |  |  |  |  |
|  | NonAbst | 16.69 ± 0.68 | 5 |  |  |  |  |  |  |
| Chronic ethanol vapor with abstinence | | | | | | | | | |
| Descriptives | | | | Statistics | | | | | |
| **Organ** | **Group** | **dCt ± SE** | **n** | **Predictor** | **Degrees of Freedom** | **Mean Square** | **F-value** | **p-value** | **Partial η2** |
| Liver | Ctrl | 9.95 ± 0.56 | 6 | Organ | 4 | 332.375 | 794.100 | 0.0000 | 0.8640 |
|  | Abst | 9.28 ± 0.28 | 6 | Group | 1 | 9.695 | 23.163 | 0.1343 | 0.0443 |
| Lung | Ctrl | 3.99 ± 0.86 | 6 | Organ*Group | 4 | 2.870 | 0.6857 | 0.6052 | 0.0520 |
|  | Abst | 4.31 ± 1.29 | 6 | Error | 50 | 4.186 |  |  |  |
| Ileum | Ctrl | 2.24 ± 0.24 | 6 |  |  |  |  |  |  |
|  | Abst | 2.16 ± 0.34 | 6 |  |  |  |  |  |  |
| Kidney | Ctrl | 10.16 ± 1.21 | 6 |  |  |  |  |  |  |
|  | Abst | 8.21 ± 1.00 | 6 |  |  |  |  |  |  |
| Heart | Ctrl | 16.47 ± 0.71 | 6 |  |  |  |  |  |  |
|  | Abst | 14.82 ± 1.02 | 6 |  |  |  |  |  |  |

**Additional analyzes of the RT-qPCR data of Ace2, Mas and Tmprss2 in the brain of ethanol vapor exposed animals**

**Suppl. Tab. 4**: Factorial ANOVA of Ace2 dCt values in the non abstinent, and abstinent models

| Chronic ethanol vapor without abstinence | | | | | | | | | |
| --- | --- | --- | --- | --- | --- | --- | --- | --- | --- |
| Descriptives | | | | Statistics | | | | | |
| **Region** | **Group** | **dCt ± SE** | **n** | **Predictor** | **Degrees of Freedom** | **Mean Square** | **F-value** | **p-value** | **Partial η2** |
| Olf.Bulb | Ctrl | 14.21 ± 0.21 | 6 | Region | 1 | 229.408 | 685.81 | 0.0000 | 0.9758 |
|  | NonAbst | 13.65 ± 0.38 | 4 | Group | 1 | 0.021 | 0.06 | 0.8060 | 0.0036 |
| PFC | Ctrl | 20.42 ± 0.16 | 6 | Region*Group | 1 | 1.297 | 3.88 | 0.0655 | 0.1857 |
|  | NonAbst | 20.86 ± 0.31 | 5 | Error | 17 | 0.335 |  |  |  |
| Chronic ethanol vapor with abstinence | | | | | | | | | |
| Descriptives | | | | Statistics | | | | | |
| **Region** | **Group** | **dCt ± SE** | **n** | **Predictor** | **Degrees of Freedom** | **Mean Square** | **F-value** | **p-value** | **Partial η2** |
| Olf.Bulb | Ctrl | 17.01 ± 1.28 | 6 | Region | 1 | 136.730 | 27.798 | 0.0000 | 0.5816 |
|  | Abst | 16.47 ± 1.25 | 6 | Group | 1 | 0.662 | 0.135 | 0.7175 | 0.0067 |
| PFC | Ctrl | 21.58 ± 0.19 | 6 | Region*Group | 1 | 0.264 | 0.054 | 0.8192 | 0.0027 |
|  | Abst | 21.45 ± 0.19 | 6 | Error | 20 | 4.919 |  |  |  |

**Suppl. Tab. 5**: Factorial ANOVA of Mas dCt values in the non abstinent, and abstinent models

| Chronic ethanol vapor without abstinence | | | | | | | | | |
| --- | --- | --- | --- | --- | --- | --- | --- | --- | --- |
| Descriptives | | | | Statistics | | | | | |
| **Region** | **Group** | **dCt ± SE** | **n** | **Predictor** | **Degrees of Freedom** | **Mean Square** | **F-value** | **p-value** | **Partial η2** |
| Olf.Bulb | Ctrl | 2.06 ± 0,14 | 6 | Region | 1 | 187.843 | 2.535.501 | 0.0000 | 0.9937 |
|  | NonAbst | 2.62 ± 0.17 | 4 | Group | 1 | 0.193 | 2.602 | 0.1263 | 0.1399 |
| PFC | Ctrl | 8.61 ± 0.06 | 5 | Region*Group | 1 | 0.628 | 8.482 | 0.0102 | 0.3465 |
|  | NonAbst | 8.45 ± 0.11 | 5 | Error | 16 | 0.074 |  |  |  |
| Chronic ethanol vapor with abstinence | | | | | | | | | |
| Descriptives | | | | Statistics | | | | | |
| **Region** | **Group** | **dCt ± SE** | **n** | **Predictor** | **Degrees of Freedom** | **Mean Square** | **F-value** | **p-value** | **Partial η2** |
| Olf.Bulb | Ctrl | 3.46 ± 0.63 | 6 | Region | 1 | 155.337 | 132.309 | 0.0000 | 0.8687 |
|  | Abst | 3.37 ± 0.60 | 6 | Group | 1 | 0.021 | 0.018 | 0.8954 | 0.0009 |
| PFC | Ctrl | 8.40 ± 0.12 | 6 | Region*Group | 1 | 0.142 | 0.121 | 0.7321 | 0.0060 |
|  | Abst | 8.61 ± 0.04 | 6 | Error | 20 | 1.174 |  |  |  |

**Suppl. Tab. 6**: Factorial ANOVA of Tmprss2 dCt values in the non abstinent, and abstinent models

| Chronic ethanol vapor without abstinence | | | | | | | | | |
| --- | --- | --- | --- | --- | --- | --- | --- | --- | --- |
| Descriptives | | | | Statistics | | | | | |
| **Region** | **Group** | **dCt ± SE** | **n** | **Predictor** | **Degrees of Freedom** | **Mean Square** | **F-value** | **p-value** | **Partial η2** |
| Olf.Bulb | Ctrl | 14.43 ± 0.29 | 6 | Region | 1 | 302.585 | 138.016 | 0.0000 | 0.8846 |
|  | NonAbst | 14.26 ± 1.04 | 5 | Group | 1 | 0.110 | 0.050 | 0.8255 | 0.0028 |
| PFC | Ctrl | 21.85 ± 0.58 | 6 | Region*Group | 1 | 0.004 | 0.002 | 0.9650 | 0.0001 |
|  | NonAbst | 21.73 ± 0.51 | 5 | Error | 18 | 2.192 |  |  |  |
| Chronic ethanol vapor with abstinence | | | | | | | | | |
| Descriptives | | | | Statistics | | | | | |
| **Region** | **Group** | **dCt ± SE** | **n** | **Predictor** | **Degrees of Freedom** | **Mean Square** | **F-value** | **p-value** | **Partial η2** |
| Olf.Bulb | Ctrl | 15.64 ± 0.91 | 6 | Region | 1 | 248.895 | 63.260 | 0.0000 | 0.7598 |
|  | Abst | 14.81 ± 0.98 | 6 | Group | 1 | 0.058 | 0.015 | 0.9042 | 0.0007 |
| PFC | Ctrl | 21.35 ± 0.83 | 6 | Region*Group | 1 | 3.255 | 0.827 | 0.3739 | 0.0397 |
|  | Abst | 21.98 ± 0.39 | 6 | Error | 20 | 3.934 |  |  |  |

**Additional analyzes in situ gene expression data of Tmprss2 and Mas on different brain regions**

**Suppl. Tab. 7**: Brain region-wise multiple t-tests of Tmprss2 nanocurie per mg (nCi/g) values in the abstinent model.

| **Brain region** | **Group** | **nCi/g ± SE** | **n** | **p-value** |
| --- | --- | --- | --- | --- |
| Cg | Ctrl | 444.75 ± 46.02 | 6 | 0.354626 |
|  | Abst | 375.06 ± 29.83 | 5 |  |
| IL | Ctrl | 430.58 ±34.46 | 6 | 0.560285 |
|  | Abst | 474.40 ± 38.54 | 5 |  |
| PrL | Ctrl | 472.17 ± 45.86 | 8 | 0.546338 |
|  | Abst | 434.69 ± 36.74 | 8 |  |
| OF | Ctrl | 375.43 ± 33.89 | 8 | 0.530227 |
|  | Abst | 335.06 ± 14.42 | 7 |  |
| CPu | Ctrl | 366.79 ± 30.70 | 8 | 0.412027 |
|  | Abst | 315.81 ± 24.05 | 8 |  |
| AcbC | Ctrl | 429.53 ± 37.53 | 8 | 0.314156 |
|  | Abst | 366.95 ± 24.80 | 8 |  |
| AcbSh | Ctrl | 464.47 ± 44.73 | 8 | 0.314156 |
|  | Abst | 365.56 ± 21.17 | 8 |  |
| PVN | Ctrl | 574.42 ± 80.13 | 7 | 0.422486 |
|  | Abst | 522.82 ± 57.56 | 8 |  |
| CA1 | Ctrl | 743.87 ± 48.87 | 8 | 0.088593 |
|  | Abst | 652.22 ± 49.62 | 8 |  |
| CA3 | Ctrl | 631.56 ± 48.80 | 8 | 0.09806 |
|  | Abst | 525.48 ± 43.73 | 8 |  |
| CA4 | Ctrl | 714.45 ± 68.96 | 8 | 0.133632 |
|  | Abst | 611.41 ± 38.98 | 8 |  |
| DG | Ctrl | 870.69 ± 58.01 | 8 | 0.133632 |
|  | Abst | 777.31 ± 65.55 | 8 |  |
| BLA | Ctrl | 463.19 ± 50.33 | 8 | 0.556342 |
|  | Abst | 426.64 ± 32.93 | 8 |  |
| CeA | Ctrl | 576.59 ± 40.80 | 8 | 0.104875 |
|  | Abst | 472.04 ± 20.42 | 7 |  |
| BMA | Ctrl | 501.18 ± 50.54 | 8 | 0.402017 |
|  | Abst | 449.09 ± 38.81 | 8 |  |
| MeA | Ctrl | 570.55 ± 60.71 | 8 | 0.547631 |
|  | Abst | 533.19 ± 39.18 | 8 |  |
| VTA | Ctrl | 343.79 ± 41.52 | 8 | 0.529691 |
|  | Abst | 382.84 ± 22.02 | 8 |  |
| NTS | Ctrl | 433.71 ± 58.94 | 7 | 0.664279 |
|  | Abst | 461.62 ± 34.20 | 8 |  |

**Suppl. Tab. 8**: Brain region-wise multiple t-tests of Mas nanocurie per mg (nCi/g) values in the abstinent model.

| **Brain region** | **Group** | **nCi/g ± SE** | **n** | **p-value** |
| --- | --- | --- | --- | --- |
| Cg | Ctrl | 1.19 ± 0.20 | 5 | 0.84391 |
|  | Abst | 0.96 ± 0.09 | 6 |  |
| IL | Ctrl | 1.79 ± 0.02 | 3 | 0.488844 |
|  | Abst | 0.77 ± 0.30 | 4 |  |
| PrL | Ctrl | 1.02 ± 0.29 | 6 | 0.776275 |
|  | Abst | 1.35 ± 0.24 | 5 |  |
| OF | Ctrl | 0.58 ± 0.22 | 7 | 0.840353 |
|  | Abst | 0.80 ± 0.25 | 5 |  |
| CA1 | Ctrl | 2.29 ± 0.22 | 8 | 0.876506 |
|  | Abst | 2.44 ± 0.41 | 8 |  |
| CA3 | Ctrl | 19.70 ± 1.15 | 8 | 0.101525 |
|  | Abst | 21.29 ± 0.92 | 8 |  |
| CA4 | Ctrl | 10.00 ± 0.58 | 8 | 0.488232 |
|  | Abst | 10.66 ± 0.48 | 8 |  |
| DG | Ctrl | 19.19 ± 1.66 | 8 | 0.465824 |
|  | Abst | 19.90 ± 1.17 | 8 |  |
| BLA | Ctrl | 1.67 ± 0.11 | 7 | 0.859646 |
|  | Abst | 1.49 ± 0.21 | 7 |  |
| CeA | Ctrl | 0.57 ± 0.12 | 7 | 0.733031 |
|  | Abst | 0.94 ± 0.42 | 6 |  |

**Suppl. Tab. 9**: Factorial ANOVA of Tmprss2 dCt values in 0.5 g/kg, 1.5 g/kg, 3.5 g/kg acute ethanol treatment.

| Acute treatment | | | | | | | | | |
| --- | --- | --- | --- | --- | --- | --- | --- | --- | --- |
| Descriptives | | | | Statistics | | | | | |
| **Organ** | **Dose** | **dCt ± SE** | **n** | **Predictor** | **Degrees of Freedom** | **Mean Square** | **F-value** | **p-value** | **Partial η2** |
| Liver | Saline | 9.19 ± 0.17 | 6 | Dose | 3 | 1.459 | 3.107 | 0.0330 | 0.1345 |
|  | 0.5 EtOH | 9.17 ± 0.25 | 6 | Organ | 2 | 476.576 | 1.015 | 0.0000 | 0.9713 |
|  | 1.5 EtOH | 9.10 ± 0.21 | 6 | Dose*Organ | 6 | 1.148 | 2,445 | 0.0351 | 0.1964 |
|  | 3.0 EtOH | 8.40 ± 0.21 | 6 | Error | 60 | 0.469 |  |  |  |
| Lung | Saline | 12.55 ± 0.54 | 6 |  |  |  |  |  |  |
|  | 0.5 EtOH | 12.02 ± 0.32 | 6 |  |  |  |  |  |  |
|  | 1.5 EtOH | 10.92 ± 0.26 | 6 |  |  |  |  |  |  |
|  | 3.0 EtOH | 11.74 ± 0.40 | 6 |  |  |  |  |  |  |
| Ileum | Saline | 2.99 ± 0.15 | 6 |  |  |  |  |  |  |
|  | 0.5 EtOH | 3.21 ± 0.18 | 6 |  |  |  |  |  |  |
|  | 1.5 EtOH | 2.91 ± 0.28 | 6 |  |  |  |  |  |  |
|  | 3.0 EtOH | 3.17 ± 0.07 | 6 |  |  |  |  |  |  |

**Supplementary Materials and Methods**

**Acute ethanol IP treatment**

For the acute Ethanol IP treatment, male Wistar rats (Envigo, Ettlingen, Germany) at the age of seven weeks were randomly distributed in groups and cages of three animals per cage. The animals were habituated to the room and the scientists for two weeks, so they were at the same age as the sub-chronic treatment group at the time point of tissue harvesting. The animals were randomly divided into four different groups, six animals per group, according to the different ethanol concentrations applied: 0.5 g/kg, 1.5 g/kg, 3.0 g/kg and a saline treated control group. The animals were decapitated 6 h after the IP injection and 2 h before the end of the active cycle (ZT22). The organs lung, liver, and ileum were dissected right after, snap frozen in isopentane and stored in -80°C until further processing. Suppl. Fig. 3 represents the overview of this experimental design.

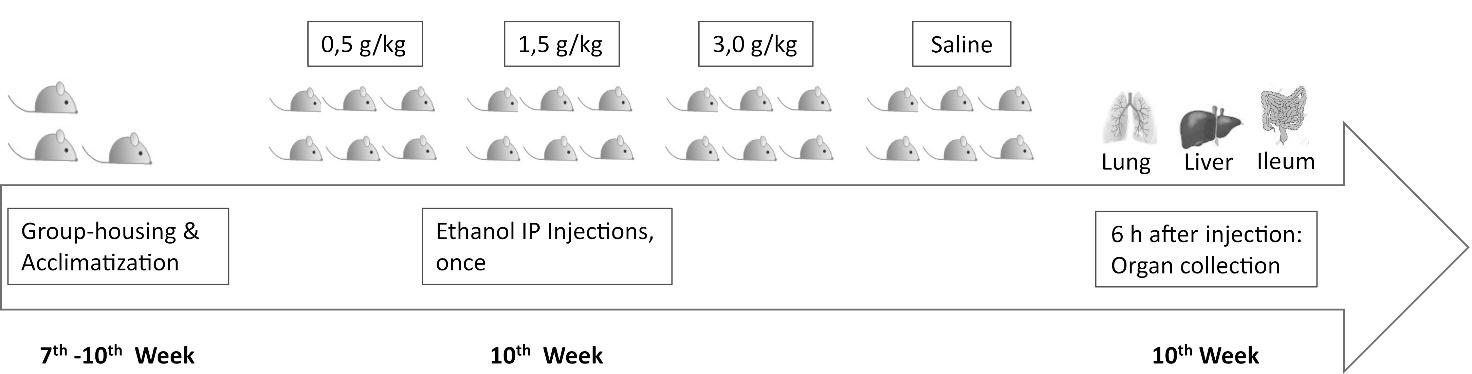

**Suppl. Fig. 4**: Time-table of acute ethanol IP injections in male Wistar rats. The animals arrived in the laboratory at the age of seven weeks. After one week of acclimatization to the housing room, one single IP injection was performed of either saline, or ethanol at 0.5 g/kg, 1.5 g/kg, 3.0 g/kg dose depending on the group. Lung, heart, and ileum were dissected 6 h after the injection.

**RT-qPCR**

**Suppl. Tab. 10:** Primer sequences of the primer pairs used for RT-qPCR

| **Species** | **Name** | **Sequence Forward** | **Sequence Reverse** | **Product length** |
| --- | --- | --- | --- | --- |
| rat | Ace2 | 5’-TGCGTATGAATGGACCGA-3’ | 5’-CAGGATAACAATGCCAACCAC-3’ | 388 |
| rat | Mas | 5’-CCCAAGCACCAGTCGGCATTC-3’ | 5’-AGGCCCATGTGTTCTTCCGTA-3’ | 229 |
| rat | Tmprss2 | 5’-GGTTTGGGCACATAGG-3’ | 5’-CACAGGCAATGGGTAGTGTTC-3’ | 251 |
| rat | Gapdh | 5′-CATGAGAAGTATGACAACAGCCT-3′ | 5′-AGTCCTTCCACGATACCAAAGT-3 | 113 |

**In-situ hybridization in rat brain tissue**

**Fixation**

10 μm coronal brain tissue sections were first fixed as described previously (Hansson et al., 2008). They were warmed to room temperature, then incubated in the following series of solutions: 4% PFA in PBS for 15 min, PBS for 10 min, sterile water for 5 min (twice), 0.1 M HCl for 10 min, PBS for 15 min (twice), 0.1 M triethanolamine (pH 8) with 0.25 % acetic anhydride for 20 min, PBS for 5 min (twice), sterile water for 1 min, graded series of ethanol (70%, 80%, 99%) for 2 min each. The so treated slices were then air dried and incubated at -80 °C in sealed boxes with silica gel to avoid moisture.

**Probe generation**

Gene-specific riboprobes for Mas and Tmprss2 were generated via a PCR based approach (Betty et al., 1998; Stoflet et al., 1988), by adding T3 (anti-sense) and T7 (sense) phage promoter sequences to the template DNA with subsequent in vitro transcription. In detail: 200 ng of DNA was incubated with 1x transcription buffer, 12.5 nmol ATP, CTP, GTP, 50 pmol UTP, and 125 pmol Uridine 5-(α-thio)triphosphate-[35S] (Perkin Elmer, Massachussets, USA), 1 U RNase inhibitor and 1 U polzmerase for 120 min at 37 °C. The DNA template was then digested by DNAseI (20 min, 37 °C) and the riboprobes were finally purified using Illustra^TM^ MicroSpin S-200 HR Colums according to the manufacturer protocol.

**Probe hybridization and washing**

Before performing the hybridization step. Three different hybridization temperatures (37, 46, 55 °C) were tested for each probe. We then selected the temperature that resulted in the best signal to noice ratio.

The fixed tissue sections were incubated in prehybridization buffer (100 mM Tris-HCl, pH 7.6; 5 mM EDTA; 5x Denhardt’s solution, 1.25 mg/ml yeast tRNA, 40 mM NaCl) diluted 1:1 with deionized formamide (2h, 37 °C), then incubated with hybridization mix (50 % deionized formamide, 150 mM DTT, 330 mM NaCl, 10% dextran sulfate, 1x basic mix (20 mM Tris-HCl, pH 7.6; 1 mM EDTA; 1x Denhardt’s solution; 0.5 mg/ml yeast tRNA; 0.1 mg/ml polyadenylic acid) containing 10 000 CPM/μl riboprobes at 37 or 55 °C for Tmprss2 and Mas overnight, respectivly. Subsequently, sections were washed twice in 1x SSC (30 min, 42 °C each), then treated with RNase (2 mg/100 ml RNase buffer) at 37 °C for 1 h, to remove unspecific signal. The enzyme reaction was finally stopped via two washing steps in 1x SSC (30 min, 55 °C).

The sections were then dipped in water for 2 min and dehydrated in a graded series of ethanol (70%, 80%, 99%; 2 min each). For signal detection, Fujifilm BAS imaging plates were exposed to the sections for 1 week.
